# Tuning of Chitosan with Lignin derived Bioactive Properties to develop a Lignin Reinforced and Sustainable Food Packaging Biomaterial

**DOI:** 10.1101/2024.05.03.592363

**Authors:** Sumona Garg, Althuri Avanthi

## Abstract

The crucial component of food storage, preservation, and transportation is food packaging. Biodegradable biopolymers are a major area of focus for the future development of food packaging materials due to their ability to mitigate adverse environmental impact by reducing plastic pollution and promoting sustainable waste management practices. Exploring renewable resources is crucial for facilitating the transition from non-renewable practices to sustainability. Chitosan is known for its superior film-forming abilities but has limited antibacterial activity. The inherent properties of lignin, including its high tensile strength, antioxidant, antimicrobial, and UV barrier ability, can contribute to enhance chitosan film performance when added as a co-polymer, making it an active material for food packaging applications. The present work explores the acid-alkali treatment to extract lignin from sugarcane tops, an abundant agricultural waste, and the application of extracted lignin in biopolymer-based hydrogel synthesis for food packaging. The goal is to enhance the hydrogel formulation by incorporation and optimisation of lignin that holds high antioxidant, antimicrobial, UV barrier, and mechanical properties along with significantly low water transmissibility. This study introduces a novel approach by utilizing lignin extracted from sugarcane tops (SCT) rather than commercially derived lignin, thereby expanding the raw material scope of lignin applications. The incorporation of higher proportions of lignin in the hydrogel formulations represents an advancement over reported studies, aimed at improving the bioactivity of the hydrogel by leveraging its advantageous characteristics emanating from lignin. This approach can also reduce the dependency on chitosan which is relatively expensive. Further, the modified synthesis of hydrogels expedited through heating method contributes to shorten the time duration needed for hydrogel film casting and drying.

**Graphical Abstract:** 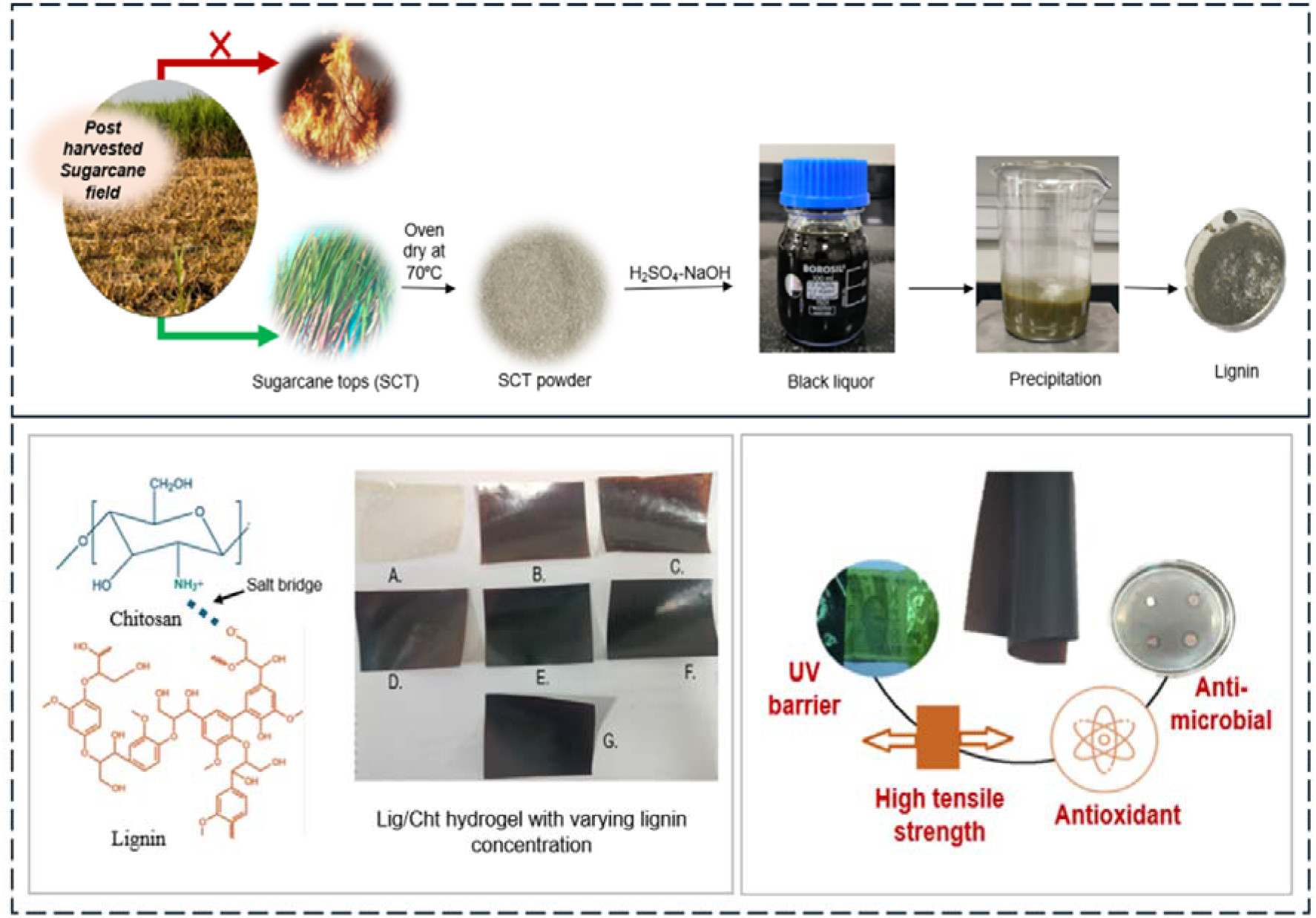

**Highlights:**

- Enhanced lignin extraction from sugarcane tops using Plackett Burman Design
- Formulating green packaging hydrogels through valorisation of sugarcane tops
- Heating-based short time casting method for Lignin/Chitosan hydrogel synthesis
- Optimization of lignin content in the hydrogels for balanced mechanical and bioactive properties

## 1.0 Introduction

The quest for sustainable and eco-friendly packaging solutions has fuelled the exploration of lignin chitosan hydrogels as a promising alternative in the realm of sustainable packaging materials. Chitosan, derived from bio-waste using energy-efficient methods, emerges as an economically viable option due to its high mechanical strength, barrier properties, and biodegradability. This green material has garnered significant attention from both the packaging and medical industries, prompting researchers to delve into novel processing techniques for chitosan hydrogels. Despite the film forming attributes of chitosan hydrogels, the challenge lies in addressing their high brittleness, fragility, and poor mechanical properties [1]. This necessitates further investigation to ascertain the sustainability of the material for packaging applications.

Around 50 million tons of lignin, the second most abundant renewable organic polymer on earth, is produced each year as a byproduct of the paper pulping industry [2]. Lignin has enormous promise for plastic applications that are environmentally sustainable. Lignin-based biopolymers have been widely accepted as an approach towards environmental sustainability in plastic usage [3]. The addition of lignin to various polysaccharide-based hydrogels has been found to significantly improve their physical and mechanical properties. For instance, incorporating lignin into chitosan hydrogels enhanced antimicrobial activity against both Gram-positive (*Bacillus subtilis, Staphylococcus aureus)* and Gram-negative (*Pseudomonas aeruginosa, Escherichia coli)* [4, 5]. A study found that incorporating lignin into chitosan films resulted in decreased water content and solubility, while enhancing tensile strength, ductility, and imparting significant antimicrobial activity against both Gram-positive and Gram-negative bacteria, suggesting potential for developing active food packaging materials .Further studies follow incorporating 30 wt% deep eutectic solvent (DES) with lignin that improved elasticity, water contact angle, and radical scavenging rate of chitosan hydrogels. Lignin also reduced water vapor permeability due to its hydrophobicity. The hydrogel with lignin exhibited significant antibacterial activity against *Staphylococcus aureus* and *Escherichia coli*, suggesting potential for bioactive food packaging [6]. Lignin nanoparticles (LNPs) produced from coconut fibre waste were incorporated into macroalgae bio-hydrogels to improve their water tolerance and mechanical characteristics. Different LNP loadings were evaluated and found to considerably improve the physical, morphological, surface roughness, structural, water resistance, mechanical, and thermal behaviour of the bio-hydrogels. The optimized bio-hydrogel exhibited hydrophobic features with a water contact angle of over 100° and high enhancement in the tensile strength of 60% [7]. The incorporation of LNP into pectin-based hydrogels also significantly enhanced their mechanical properties, hydrophobicity, and water barrier properties. The tight entanglement and strong positive interaction between pectin and LNP promote the formation of dense structures in the hydrogel, resulting in a nano-scale rough surface. Pectin-LNP composite hydrogels almost completely shield the UVB, UVC, and most of the UVA spectrum, demonstrating their strong anti-ultraviolet performance. Also, the addition of LNP significantly enhances the DPPH radical scavenging ability and antibacterial ability of the pectin hydrogel, making it a promising active packaging material for food preservation [8]. In another reported study, it was found that lignin addition improved the light barrier properties of alginate hydrogel, and FTIR analysis suggested hydrogen bonding between lignin and alginate hydrogel [9]. A reported study demonstrated that the incorporation of lignin substantially enhanced the physical and mechanical properties of alginate hydrogels, positioning them as a promising alternative material for mitigating lipid oxidation in food systems [10]. On the contrary, it is important to note that a decrease in tensile strength with tobacco lignin addition to polypropylene (PP) was observed which was attributed to the formation of larger lignin aggregates at higher concentrations. This led to decreased mechanical characteristics, affecting uniform distribution within the PP matrix. Therefore, optimizing lignin concentration is vital to maintain mechanical integrity and ensure uniform distribution within the polymer matrix, thereby preserving material strength and performance [11].

Sugarcane tops, a widely spread agricultural waste, possess significant potential as a sustainable feedstock for lignin extraction. The main biochemical components include cellulose (36-40% w/w), hemicellulose (28-32% w/w), Lignin (17-26% w/w), and others include ash, pectin, proteins, free sugars etc [12]. Apart from their higher lignin content, sugarcane tops offer additional advantages such as rapid growth, widespread availability, and no competition with food supply. Around 7–12 tons of SCTs can be produced from 1 ha of the sugarcane field [13]. On large farms the tops are burned off before the cane is processed for disposal, while on small farms the tops are cut for livestock feed or end up in landfills. Burning of SCTs leads to air pollution and burial is responsible for GHG emissions. Harnessing sugarcane tops as a sustainable feedstock for lignin extraction offers a dual benefit i.e., mitigating waste management issues associated with burning or landfilling while providing a valuable resource for lignin-based applications.

Considering the advancements and challenges, this research aims to explore the feasibility and potential of lignin chitosan hydrogels as a sustainable antioxidant, antimicrobial and UV shielding food packaging material with low water transmissibility. The objective is to synthesize a superior lignin-chitosan (Lig/Cht) film with high mechanical strength while retaining its bio-active properties derived from lignin. This study can unleash innovation for an eco-friendly packaging solution with enhanced functionality and sustainability.

## 2.0 Materials and methods

### 2.1 Materials

Sugarcane tops was procured from the local fields of Kandi, Sangareddy, Telangana, India. Feedstock was pulverised and sieved to obtain particles of mesh size 0.25mm and 1mm, and oven dried overnight at 60 °C. Chitosan (Low MW) extrapure, 10-150m.Pas, 90% deacetylation degree and lactic acid were purchased from Sisco Research Laboratories Pvt. Ltd. (SRL), India. *Lactobacillus sakei* and *Bacillus subtilis* with NCIM Accession No 5645 and 2439 were purchased from National Collection of Industrial Microorganisms (NCIM), Pune, India. Sodium hydroxide, MRS media, and nutrient broth, and agar media were obtained from SRL, India. Sulfuric acid is procured from SD Fine chemical.

### 2.2 Characterization of SCT biomass

The lignin composition of SCT was estimated using the standard NREL protocol based on gravimetry.

### 2.3 Extraction of lignin from sugarcane tops (SCT)

Sugarcane tops (SCT) dry powdered sample (5-10g) was mixed with 100ml of 1%-2% H_2_SO_4_ (v/v) followed by autoclaving at 121°C at 15psi for 10-15 mins. The solid was dried at 50°C overnight and treated with 3-5 % of 100ml NaOH treatment and autoclaved at 121°C at 15psi for 10-15 mins. The alkali lignin black liquor obtained after vacuum filtration was precipitated by addition of concentrated HCl (pH ∼2) and centrifugated at 5000 rpm for 15 min. The precipitate was washed with distilled water till neutral pH and dried in an oven set at 50 °C [14].

### 2.4 Experimental design and statistical analysis for enhanced lignin recovery

The Plackett-Burman (PB) experimental design was employed using Minitab 17 software to explore the key factors influencing lignin yield from SCT using the acid-alkali method. A total of 12 experimental runs were carried out (Table 1), each triplicated, encompassing various combinations of factors at two levels: high (+) and low (-). These factors included feedstock concentration (5%,10% (w/v)), sulfuric acid (HLSOL) concentration (1%,2% (v/v)), sodium hydroxide (NaOH) concentration (3%,5% (w/v)), reaction time (10min, 15min), and particle size (0.25mm, 1mm). The resulting lignin yields from these experimental runs underwent statistical analysis, using analysis of variance (ANOVA) to ascertain the significance of each factor and the effect of their potential interactions on lignin extraction efficiency. Additionally, graphical tools such as half normal plots and pareto charts were utilized to visually portray the relative importance of these factors and pinpoint those exerting notable influence [15].

**Table 1.**
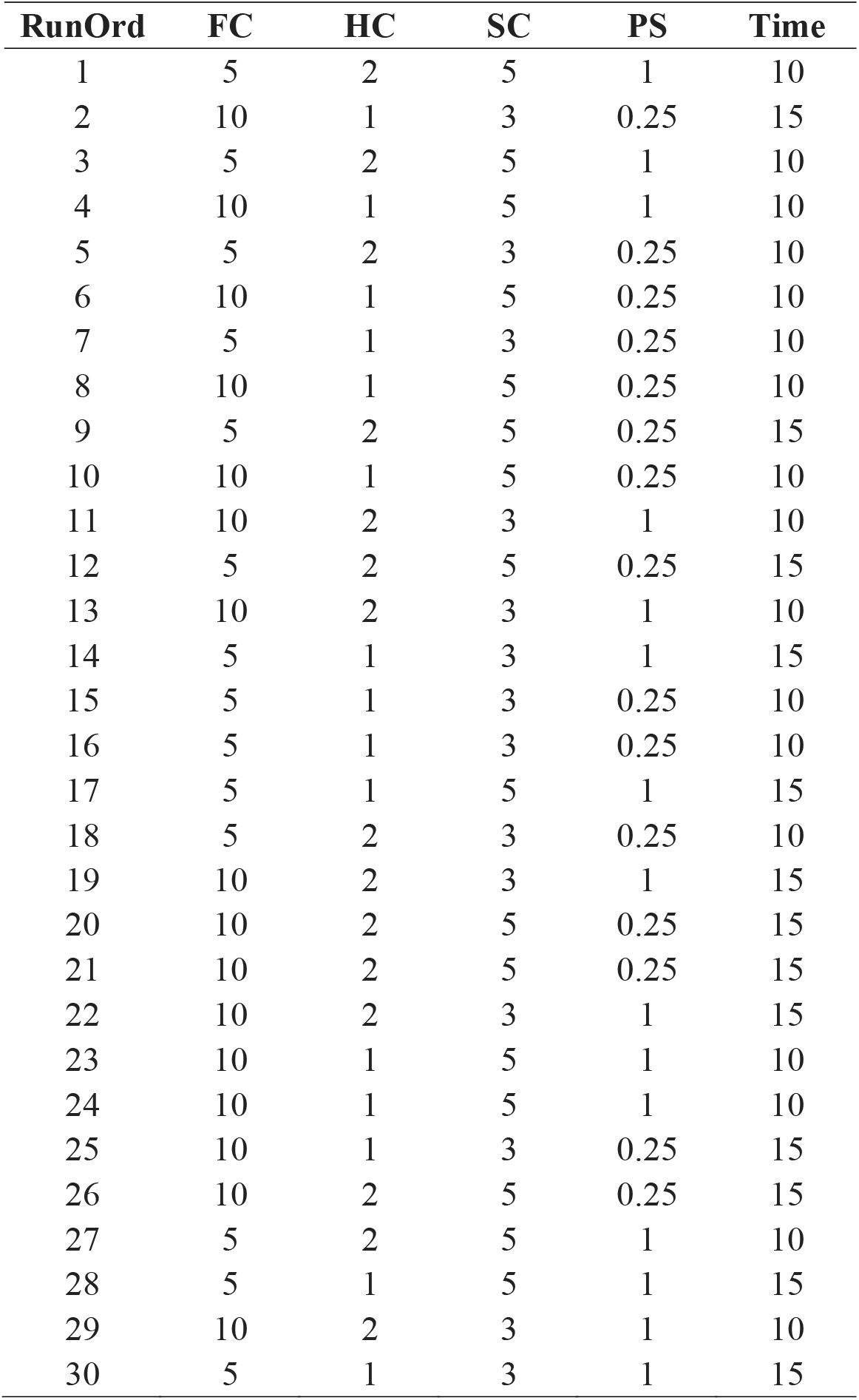
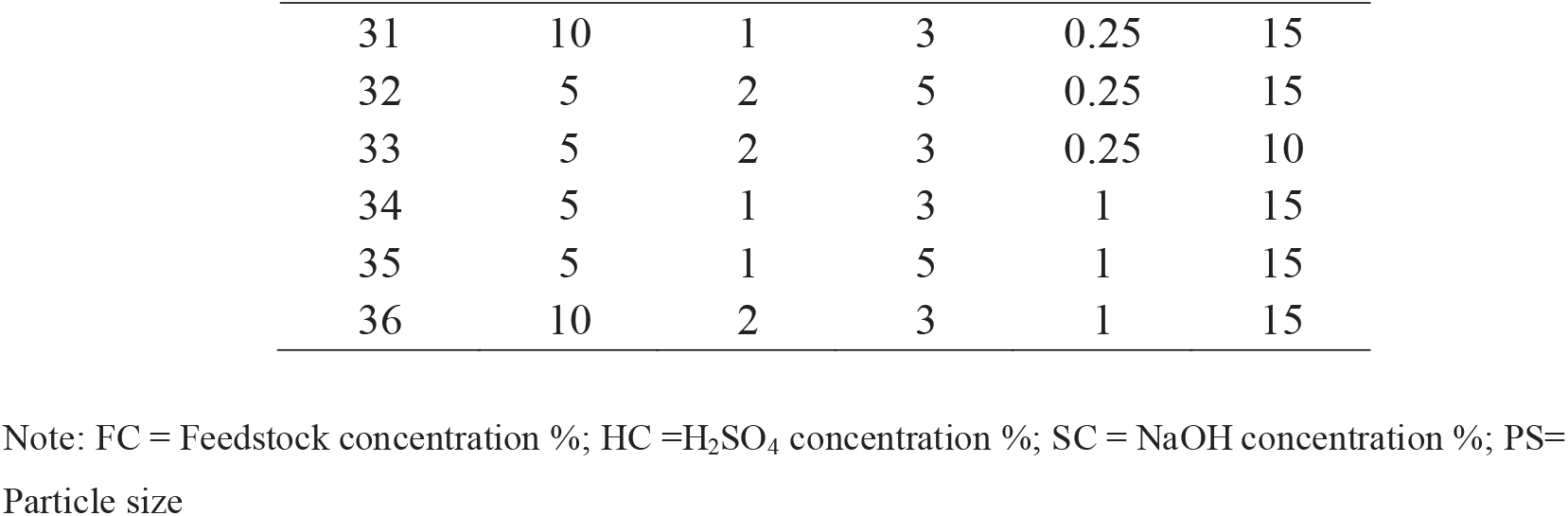
Design of experiment for optimization of lignin yield (%w/w) from sugarcane tops (SCT) based on Plackett-Burman (PB) design.

| RunOrd | FC | HC | SC | PS | Time |
| --- | --- | --- | --- | --- | --- |
| 1 | 5 | 2 | 5 | 1 | 10 |
| 2 | 10 | 1 | 3 | 0.25 | 15 |
| 3 | 5 | 2 | 5 | 1 | 10 |
| 4 | 10 | 1 | 5 | 1 | 10 |
| 5 | 5 | 2 | 3 | 0.25 | 10 |
| 6 | 10 | 1 | 5 | 0.25 | 10 |
| 7 | 5 | 1 | 3 | 0.25 | 10 |
| 8 | 10 | 1 | 5 | 0.25 | 10 |
| 9 | 5 | 2 | 5 | 0.25 | 15 |
| 10 | 10 | 1 | 5 | 0.25 | 10 |
| 11 | 10 | 2 | 3 | 1 | 10 |
| 12 | 5 | 2 | 5 | 0.25 | 15 |
| 13 | 10 | 2 | 3 | 1 | 10 |
| 14 | 5 | 1 | 3 | 1 | 15 |
| 15 | 5 | 1 | 3 | 0.25 | 10 |
| 16 | 5 | 1 | 3 | 0.25 | 10 |
| 17 | 5 | 1 | 5 | 1 | 15 |
| 18 | 5 | 2 | 3 | 0.25 | 10 |
| 19 | 10 | 2 | 3 | 1 | 15 |
| 20 | 10 | 2 | 5 | 0.25 | 15 |
| 21 | 10 | 2 | 5 | 0.25 | 15 |
| 22 | 10 | 2 | 3 | 1 | 15 |
| 23 | 10 | 1 | 5 | 1 | 10 |
| 24 | 10 | 1 | 5 | 1 | 10 |
| 25 | 10 | 1 | 3 | 0.25 | 15 |
| 26 | 10 | 2 | 5 | 0.25 | 15 |
| 27 | 5 | 2 | 5 | 1 | 10 |
| 28 | 5 | 1 | 5 | 1 | 15 |
| 29 | 10 | 2 | 3 | 1 | 10 |
| 30 | 5 | 1 | 3 | 1 | 15 |
| 31 | 10 | 1 | 3 | 0.25 | 15 |
| 32 | 5 | 2 | 5 | 0.25 | 15 |
| 33 | 5 | 2 | 3 | 0.25 | 10 |
| 34 | 5 | 1 | 3 | 1 | 15 |
| 35 | 5 | 1 | 5 | 1 | 15 |
| 36 | 10 | 2 | 3 | 1 | 15 |

Note: FC = Feedstock concentration %; HC =H<sub>2</sub>SO<sub>4</sub> concentration %; SC = NaOH concentration %; PS=
Particle size

### 2.5 Preparation of hydrogel casting solutions

To expedite the hydrogel setting process, a modified protocol was adopted for preparing the chitosan solution, incorporating a novel heating method. This modification aims to reduce the time required for hydrogel formation. To the best of our knowledge, no previous studies have reported such an approach. To obtain 2% (w/v) chitosan solution, 2 g chitosan powder was dispersed into 1% (v/v) lactic acid solution followed by stirring at 400 rpm until transparent solution is formed. The extracted alkali lignin is dissolved in 70% (v/v) ethanol and stirred at 60 °C for 24 h and vacuum filtrated to remove the insoluble part. Solution of different lignin concentration was prepared using 70% (v/v) ethanol such that the concentration of lignin varies from 0.5-1% (w/v). Later, 100ml of the lignin solution was added to 150ml of chitosan solution. This solution was stirred at 60 °C under 250 rpm till about 70% of the solution was evaporated [16].

### 2.6 Casting of Lig/Cht hydrogels

Teflon sheets were cut into 20 x 12 cm and placed onto a casting tray. Hydrogel casting solution (75-80 ml) was poured onto the teflon sheets and dried at 40°C in an oven for 8 h. Then, the hydrogels were peeled off from the teflon sheets and stored in zip lock bags at room temperature for further studies. All hydrogels were pre-conditioned at 25 ± 1 ◦C and 50 ± 1% relative humidity (RH) for at least 72 h before further tests. The hydrogels casted are denoted as 0.5Lig/3Cht, 0.6Lig/3Cht, 0.7Lig/3Cht, 0.8Lig/3Cht, 0.9Lig/3Cht, 1Lig/3Cht based on the individual polymer concentrations in the hydrogel formulations [16].

### 2.7 Characterization

#### 2.7.1 Fourier transform infrared spectroscopy analysis (FT-IR)

FTIR spectra of the extracted alkali lignin and commercial alkali lignin (sigma grade) were obtained using a Bruker TENSOR 37 FTIR spectrometer following standard ASTM E1252. The IR spectrum was recorded in the wavenumber range of 550–4000 cm^−1^. In this method, a transparent circular disk containing powdered oxide catalyst of spectroscopic grade, KBr, is made using a hydraulic press. The sample was mixed with KBr powder, pressed into a pellet, and then analysed using the FTIR spectrometer.

#### 2.7.3 UV barrier property

The UV Barrier property of the Lig/Cht hydrogels was analyzed using a Lambda 365 UV-vis spectrophotometer. For the measurement, hydrogel samples were cut into the size of cuvettes and the reference was taken as air. The hydrogels were scanned from 200 to 800 nm [17].

#### 2.7.4 Antioxidant activity

Hydrogel (200 mg) was mixed with 10 ml of 95% (v/v) ethanol, kept in the dark at 50°C for 3 h. Supernatant (1.5 ml) obtained after incubation was added to 1.5 ml of 0.06 mM DPPH (2,2-Diphenyl-1-picrylhydrazyl) in 95%(v/v) ethanol solution. Later, the mixture was stirred for 30 min at room temperature in darkness. The absorbance of the mixture solution was measured at 517 nm using a Jenway UV/Vis spectrophotometer. The control sample was similarly prepared using 95%(v/v) ethanol without hydrogel sample. The test was carried out in triplicates [18].

#### 2.7.5 Antimicrobial activity

The antimicrobial activity of hydrogels was screened against *Lactobacillus sakei* and *Bacillus subtilis* using the disk diffusion method. Hydrogel samples with a diameter of 1 cm were exposed to ultraviolet light for 15 min and then placed onto agar plates containing 10µl of broth inoculum with an optical density of 0.7. These plates were then incubated at 25°C for 24 h, and any microbial growth inhibition zones were measured and recorded [18].

#### 2.7.6 Tensile strength

The stress-strain curve of the hydrogels was assessed using Instron mechanical testing machine according to standard ASTM D638. The hydrogels were cut into rectangular specimens with dimensions of 20mm x 20mm and fixed on the grips of the machine. The initial grip and crosshead speed were 10 mm and 10 mm/min, respectively. The thickness of hydrogel was measured using a digital micrometre.

#### 2.7.7 Water Vapor Transmissibility Rate (WVTR)

To determine WVTR of the prepared hydrogels, 150ml conical flask was filled with 10 mL of distilled water. The mouth of the flask was then sealed with a film sample measuring 3 × 3 cm² and further secured with parafilm around the conical rim. The initial weight of the test bottle was recorded as W_1_, and it was placed in a hot air oven set at a temperature of 40 ± 2 °C for 24 h. Subsequently, the test bottles were removed from the oven, allowed to cool, and weighed again to obtain the final weight W_2_. The WVTR of the films was then calculated using the following equation (Eq. 1).

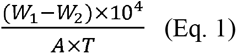

Where, T = 24 h and *A* = area of the mouth of the bottle (mm) [19].

## 3.0 Results and Discussion

### 3.1 Regression analysis for improved lignin yield from SCT

The lignin yield from sugarcane tops (SCT) was found to be significantly influenced by the reaction conditions of the acid-alkali process. For this study, Plackett-Burman derived design of experiments (DOE) was utilized, involving five factors with two levels each to optimize lignin yield. Table 2 shows lignin yield (%w/w) based on Plackett-Burman Design.

**Table 2.** Lignin yield (%w/w) based on Plackett-Burman Experimental Design.

| RunOrd | FC | HC | SC | PS | Time<br>(min) | Experimental<br>Lignin Yield<br>(%w/w) | Predicted<br>Lignin yield<br>(%w/w) |
| --- | --- | --- | --- | --- | --- | --- | --- |
| 1 | 5 | 2 | 5 | 1 | 10 | 11.45 | 12.67 |
| 2 | 10 | 1 | 3 | 0.25 | 15 | 54.02 | 47.89 |
| 3 | 5 | 2 | 5 | 1 | 10 | 11.45 | 12.67 |
| 4 | 10 | 1 | 5 | 1 | 10 | 51.64 | 41.21 |
| 5 | 5 | 2 | 3 | 0.25 | 10 | 16.52 | 17.02 |
| 6 | 10 | 1 | 5 | 0.25 | 10 | 51.18 | 48.77 |
| 7 | 5 | 1 | 3 | 0.25 | 10 | 24.27 | 23.46 |
| 8 | 10 | 1 | 5 | 0.25 | 10 | 28.56 | 48.77 |
| 9 | 5 | 2 | 5 | 0.25 | 15 | 30.41 | 22.55 |
| 10 | 10 | 1 | 5 | 0.25 | 10 | 64.99 | 48.77 |
| 11 | 10 | 2 | 3 | 1 | 10 | 26.88 | 31.58 |
| 12 | 5 | 2 | 5 | 0.25 | 15 | 32.88 | 22.55 |
| 13 | 10 | 2 | 3 | 1 | 10 | 27.14 | 31.58 |
| 14 | 5 | 1 | 3 | 1 | 15 | 22.14 | 18.23 |
| 15 | 5 | 1 | 3 | 0.25 | 10 | 28.67 | 23.46 |
| 16 | 5 | 1 | 3 | 0.25 | 10 | 14.5 | 23.46 |
| 17 | 5 | 1 | 5 | 1 | 15 | 22.15 | 21.43 |
| 18 | 5 | 2 | 3 | 0.25 | 10 | 15.88 | 17.02 |
| 19 | 10 | 2 | 3 | 1 | 15 | 28.17 | 33.90 |
| 20 | 10 | 2 | 5 | 0.25 | 15 | 47.47 | 44.66 |
| 21 | 10 | 2 | 5 | 0.25 | 15 | 24.52 | 44.66 |
| 22 | 10 | 2 | 3 | 1 | 15 | 53.45 | 33.90 |
| 23 | 10 | 1 | 5 | 1 | 10 | 27.19 | 41.21 |
| 24 | 10 | 1 | 5 | 1 | 10 | 54.14 | 41.21 |
| 25 | 10 | 1 | 3 | 0.25 | 15 | 58.66 | 47.89 |
| 26 | 10 | 2 | 5 | 0.25 | 15 | 49.50 | 44.66 |
| 27 | 5 | 2 | 5 | 1 | 10 | 26.59 | 12.67 |
| 28 | 5 | 1 | 5 | 1 | 15 | 12.67 | 21.43 |
| 29 | 10 | 2 | 3 | 1 | 10 | 28.04 | 31.58 |
| 30 | 5 | 1 | 3 | 1 | 15 | 22.65 | 18.23 |
| 31 | 10 | 1 | 3 | 0.25 | 15 | 41.33 | 47.89 |
| 32 | 5 | 2 | 5 | 0.25 | 15 | 14.61 | 22.55 |
| 33 | 5 | 2 | 3 | 0.25 | 10 | 15.02 | 17.02 |
| 34 | 5 | 1 | 3 | 1 | 15 | 11.72 | 18.23 |
| 35 | 5 | 1 | 5 | 1 | 15 | 12.46 | 21.43 |
| 36 | 10 | 2 | 3 | 1 | 15 | 27.13 | 33.90 |

Note: FC = Feedstock concentration %; HC =H<sub>2</sub>SO<sub>4</sub> concentration %; SC = NaOH concentration %; PS= Particle size

Based on the data presented in Table 1, a regression equation was developed to forecast the lignin yield percentage. The equation incorporates factors A, B, C, D, and E, which denote the concentrations of feedstock, sulfuric acid (H_2_SO_4_), sodium hydroxide (NaOH), particle size, and reaction time, respectively (Eq. 2).

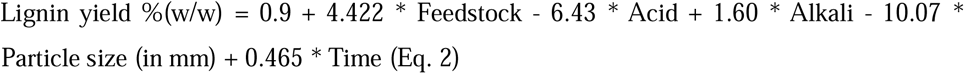

From equation 2 it can be observed that the variables namely, feedstock concentration, alkali concentration and time show direct proportional impact on lignin yield percentage whereas acid and particle size are inversely proportional to lignin yield. The adjusted R-squared and predicted R-squared values were reasonable (57.81% and 47.92%, respectively) highlighting the significant impact of the model. ANOVA for lignin yield is summarized in Table 3

**Table 3.** Analysis of variance (ANOVA) based on PB data.

| Source | DF<sup>a</sup> | Adj SS<sup>b</sup> | Adj MS<sup>c</sup> | F-Value | P-Value |
| --- | --- | --- | --- | --- | --- |
| Model | 5 | 5426.31 | 1085.26 | 10.59 | 0.000 |
| Linear | 5 | 5426.31 | 1085.26 | 10.59 | 0.000 |
| Feedstock | 1 | 4399.28 | 4399.28 | 42.93 | 0.000 |
| Acid | 1 | 372.60 | 372.60 | 3.64 | 0.066 |
| Alkali | 1 | 92.29 | 92.29 | 0.90 | 0.350 |
| Particle size | 1 | 513.57 | 513.57 | 5.01 | 0.033 |
| Time | 1 | 48.57 | 48.57 | 0.47 | 0.496 |
| Error | 30 | 3074.48 | 102.48 |  |  |
| Lack-of-Fit | 6 | 371.19 | 61.87 | 0.55 | 0.766 |
| Pure Error | 24 | 2703.28 | 112.64 |  |  |
| Total | 35 | 8500.79 |  |  |  |

<sup>a</sup>Degrees of freedom; <sup>b</sup>Sum of squares; <sup>c</sup>Mean squares.

The graph of Half Normal Plot (Fig. 1.) of the standardized effects for the response ‘Lignin yield %(w/w)’ shows the absolute standardized effects of different variables under study on lignin yield percentage. The factors represented in the plot are A=Acid (H_2_SO_4_) concentration, B= Alkali (NaOH) concentration, C=Feedstock concentration, D=Particle size in mm, and F=Time. The legend indicates significant effects are marked with a red square, while insignificant effects are marked with a blue square. The plot suggests that the ‘Feedstock’ and “Particle size” variables (factors) have a significant corelation with lignin yield as compared to acid (H_2_SO_4_), alkali (NaOH), and time that have smaller absolute standardized effects and are deemed not significant based on the analysis.

**Fig. 1.**
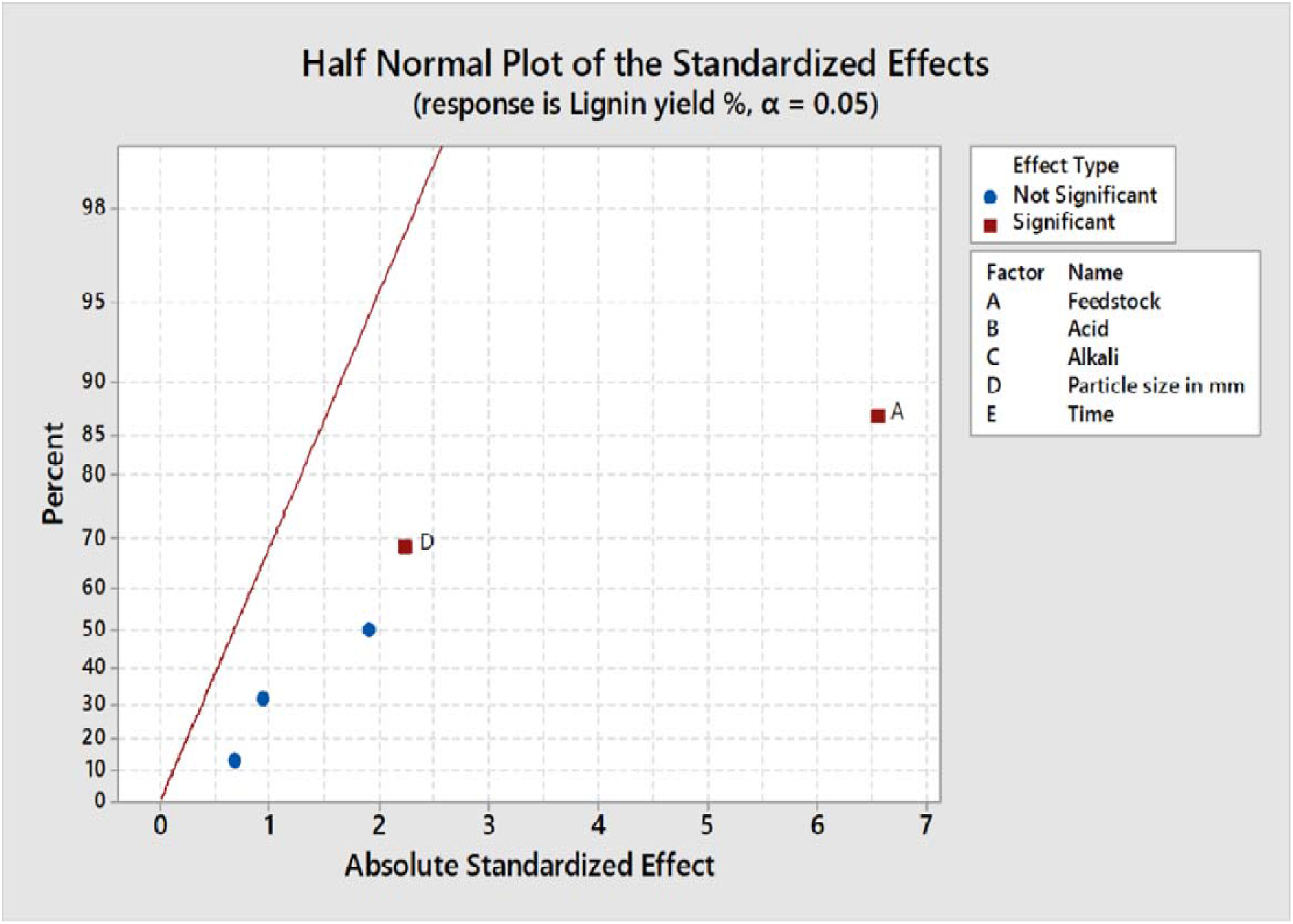
Half normal plot of the standardized effects.

The Pareto Chart (Fig. 2.) also displays the standardized effects of these factors on the lignin yield percentage, with Factor A (Feedstock concentration) and Factor B (Particle size) having the major effect. Particle size reduction enhances the lignocellulosic biomass accessibility, reactivity, and process efficiency, potentially leading to higher lignin recovery [21]. Owing this reason 0.25 mm particle size was selected for the subsequent studies. Factor E (Time) showed insignificant effect on the lignin yield which is in coherence with the findings of a previous study [22]. The outcomes from the Pareto Chart align with those from the Half Normal Plot, reinforcing the importance of ‘feedstock’ and ‘particle size’ in optimizing lignin yield. The chart is structured in a descending order of the standardized effects and indicates that the significance level (α) is 0.05. Thus, understanding these nuances in the factors impacting lignin yield is essential for refining biomass processing techniques and achieving efficient lignin extraction for various applications. Based on these results, the lignin extraction for subsequent studies was done using a concentration of 10%(w/w) feedstock with 0.25mm particle size, 1%(v/v) H_2_SO_4_, 3%(w/v) NaOH for 15 min.

**Fig. 2.**
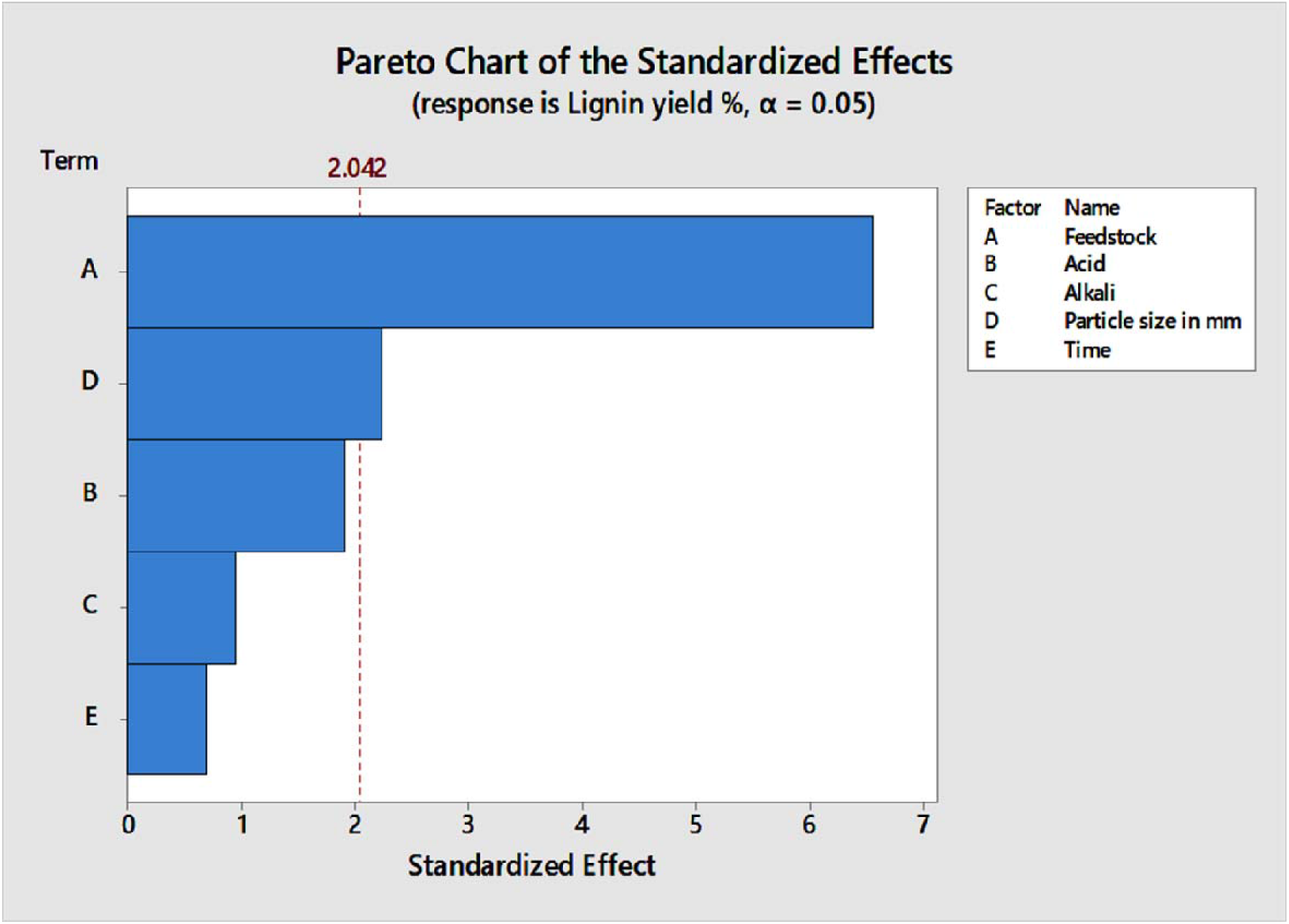
Pareto chart of the standardized effects.

Under these selected reaction conditions, the experimental lignin yield was found to be 51.33 ± 7.32% (w/w).

### 3.2 Comparative FTIR analysis of extracted lignin and commercial lignin

The FTIR analysis (Fig. 3.) of the extracted alkali lignin residue revealed characteristic peaks at 3423 cm^1^ (OH stretching) [23], 2927 cm^1^ (methyl and methylene groups) [24], 2856 cm^1^ (methyl group) [25], 1589 cm^1^ (carboxylate absorption[26]), 1510 cm^1^ (aromatic C=C bending coupled with C-H in-plane deformations) [27, 28], and 1457 cm^1^ (methoxyl C-H deformation and aromatic ring vibration) [29]. Additionally, absorption peaks at 1126 cm^1^ indicated a typical lignin structure with p-hydroxyphenylpropane (H) units [29], while C-O bands at 1035 cm^1^ were associated with secondary and primary alcohols, respectively [30]. Therefore, the FTIR data indicates that the lignin extracted from sugarcane tops (SCTs) via acid-alkali process exhibits characteristic peaks consistent with commercial grade alkali lignin. The phenolic hydroxyl groups in lignin are known for their antimicrobial and antioxidant activities, as they can disrupt microbial cell membranes and scavenge free radicals [31]. Furthermore, the aromatic structure of lignin provides UV-blocking capabilities by absorbing and dissipating UV radiation [32]. These properties make lignin extracted from SCTs a promising candidate for various applications in antimicrobial, antioxidant, and UV-protective materials.

**Fig. 3.**
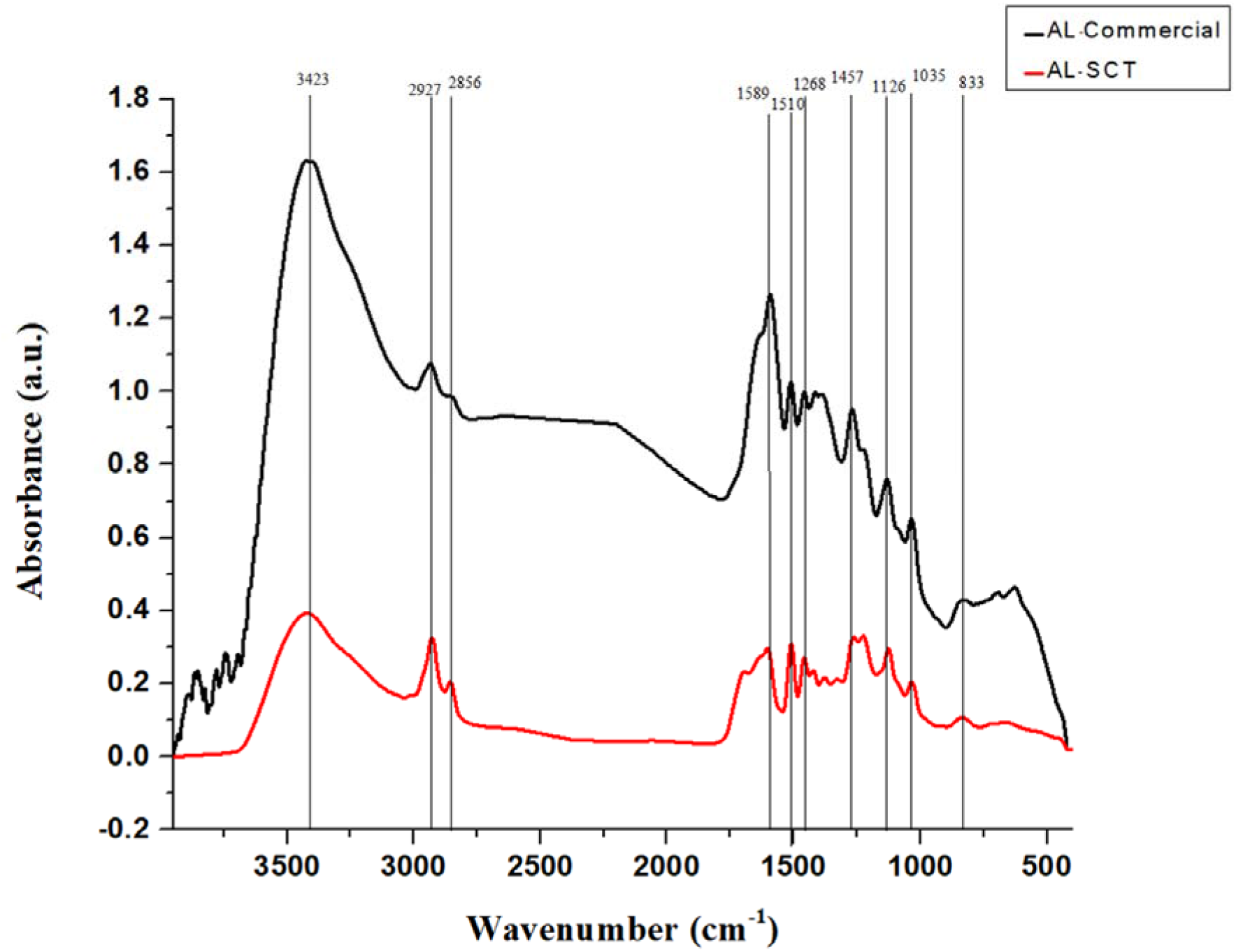
FTIR data of extracted alkali lignin and commercial lignin.

### 3.3 Influence of Lignin Concentration on Hydrogel Characteristics

#### 3.3.1 UV Barrier Property Analysis

Fig. 4. illustrates the transmittance as a function of wavelength for pure chitosan hydrogels (control, denoted as 3Cht) and hydrogels supplemented with lignin (0.5Lig/3Cht, 0.6Lig/3Cht, 0.8Lig/3Cht, and 0.9Lig/3Cht). It is imperative for hydrogels intended for food packaging applications to exhibit effective light barrier properties, particularly against ultraviolet (UV) radiation. The transmittance values for 3Cht (control) and 0.5Lig/3Cht hydrogels were 57.56% and 0.03%, respectively, across 200–4000 nm wavelength. Specifically, 3Cht (control) hydrogels demonstrated transmittance of 57.56% in the UVA region, 4.04% in the UVB region, and 0.005% in the UVC region, indicating absorption capabilities in the UVB and UVC spectra. Conversely, transmittance decreased with increasing lignin concentration, reaching 0.03% and 0.008% for UVA, 0.002% and 0.002% for UVB, and 0.005% and 0.005% for UVC in 0.5Lig/3Cht and 0.9Lig/3Cht hydrogels, respectively. The transmittance of hydrogels with higher lignin concentrations approached zero in the UV range. Similar findings were reported with PLA-grafted LNP (lignin nanoparticles) films where a significant decrease in optical transmittance in the UV range, declining from 58.7 ± 3.0% to 1.10 ± 0.01% was reported [33]. Another study revealed that lignin-containing cationic wood nanofiber (CWNF) films exhibited excellent UV absorption properties, with transmittance <1% below 380 nm [34]. Thus, in addition to lignin’s role as a reinforcing polymer, it also contributes to the enhanced bioactive properties of the hydrogel. This UV-blocking property of lignin can be attributed to its structural features containing chromophore functional groups capable of absorbing a broad spectrum of UV light [36]. These functional groups include phenolic, ketone, chromophoric, and auxochrome structures [36].

**Fig. 4.**
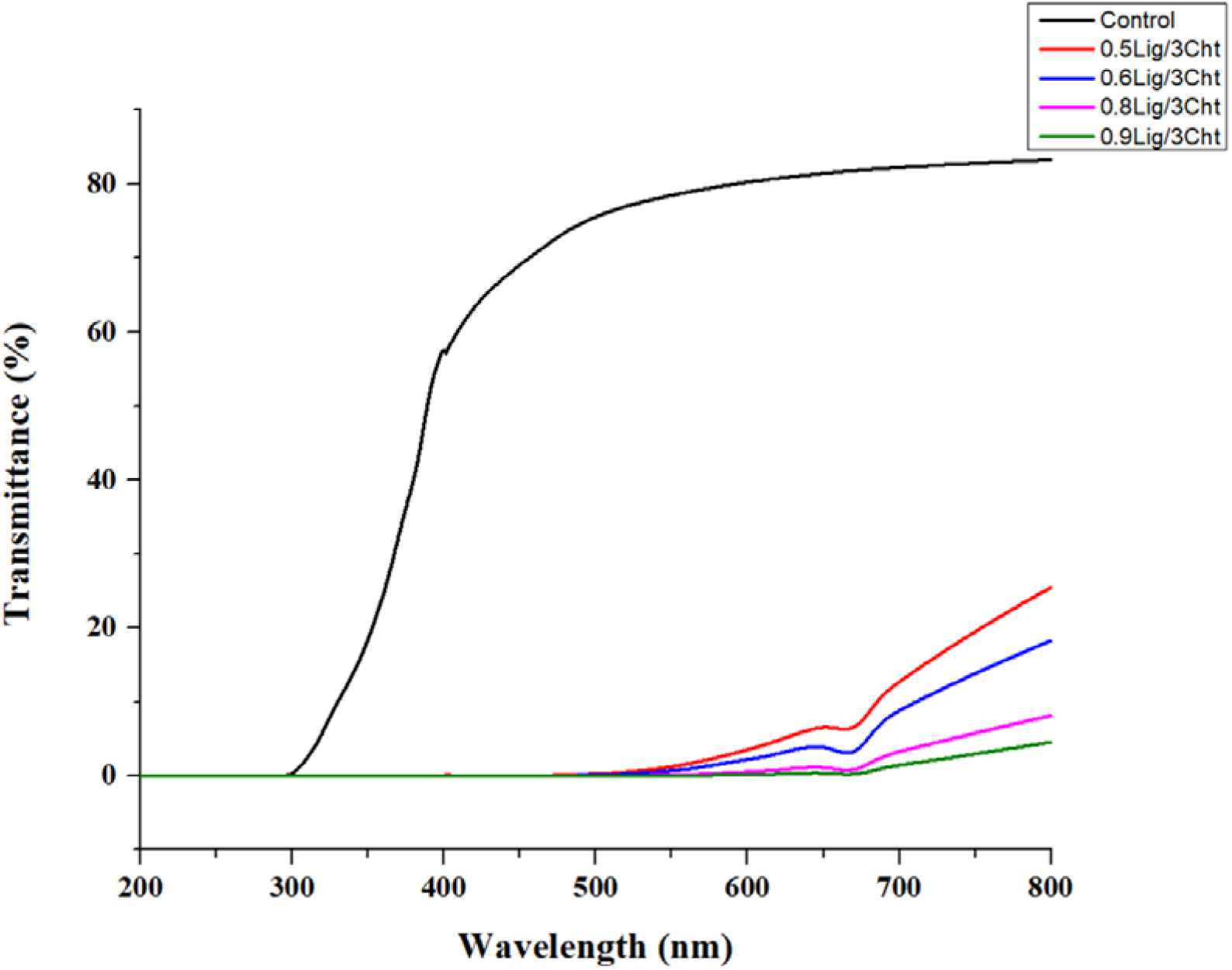
UV spectra showing UVA, UVB, and UVC blocking activity of Lig/Cht hydrogels.

#### 3.3.2 Antioxidant Property

Oxidation stands as a major factor in the deterioration of food products, undermining their nutritional and commercial worth. To combat this issue and extend the shelf-life of food items, the concept of antioxidant packaging emerges as a promising solution. Antioxidant packaging involves incorporating natural bioactive compounds, such as antioxidants, into the packaging material to inhibit oxidation and extend the shelf life of food products [34]. The utilization of agricultural waste as a source of biopolymer and additives in the development of biodegradable packaging films has gained significant interest due to the relative abundance and low cost of these waste material [35]. Moreover, lignin from these wastes can act as a potential antioxidant agent and thus enhance the performance of hydrogels. The antioxidant activity of the synthesised hydrogels was assessed using DPPH radical scavenging assay. This analysis was done to determine the effectiveness of Lig/Cht hydrogels to withstand oxidation at different lignin concentrations. The Lig/Cht hydrogel photographs are attached in the supplementary material (Fig. S1.) The control (Cht) hydrogel exhibited antioxidant activity of 7.32 ± 0.70 %. This was inextricably related to the interaction of free radicals with the hydroxyl and amino groups of chitosan. Lig/Cht hydrogels showed considerable differences in their ability to scavenge free radicals as compared to Cht film. All Lig/Cht hydrogels displayed a concentration-dependent increase in DPPH scavenging values as shown in Fig. 5. The antioxidant activity increased progressively from 31.45 ± 4.72 % to 73.68 ± 3.58 % for the 0.5Lig/3Cht to 1Lig/3Cht hydrogel, respectively. The 0.7Lig/3Cht hydrogels exhibited 57.33 ± 5.85 % activity, while the 0.8Lig/3Cht hydrogels demonstrated 70.23 ± 2.10 % activity. However, further increments in lignin concentration resulted in only marginal increases in antioxidant activity, reaching 71.42 ± 3.88 % for the 0.9Lig/3Cht and 73.68 ± 3.58 % for the 1Lig/3Cht hydrogels. The observed phenomenon could be attributed to lignin agglomeration, which may hinder the exposure of hydroxyl groups. Therefore, the 0.8Lig/3Cht hydrogel formulation represents an optimum blend for applications requiring effective antioxidant properties. Overall, this study substantiates that lignin acts as an antioxidant agent which can be accounted to its ability to donate hydrogen ions and scavenge free radicals. Its antioxidant activity is primarily attributed to the phenolic hydroxyl structure, with the hydroxyl group playing a central role in this process [36]. Thus, the free radical scavenging capacity of these Lig/Cht hydrogels can be exploited for antioxidant packaging of food materials. A study reported the antioxidant capacity of neat PLA, LNPs and PLA-grafted LNPs films. It was found that compared to neat PLA, the antioxidant capacity increased by over 10 times in LNPs and 12 times in PLA-grafted LNP films. This significant enhancement underscores the potential of lignin-based hydrogels for food packaging applications, where the preservation of food quality and shelf life is crucial [37].

**Fig. 5.**
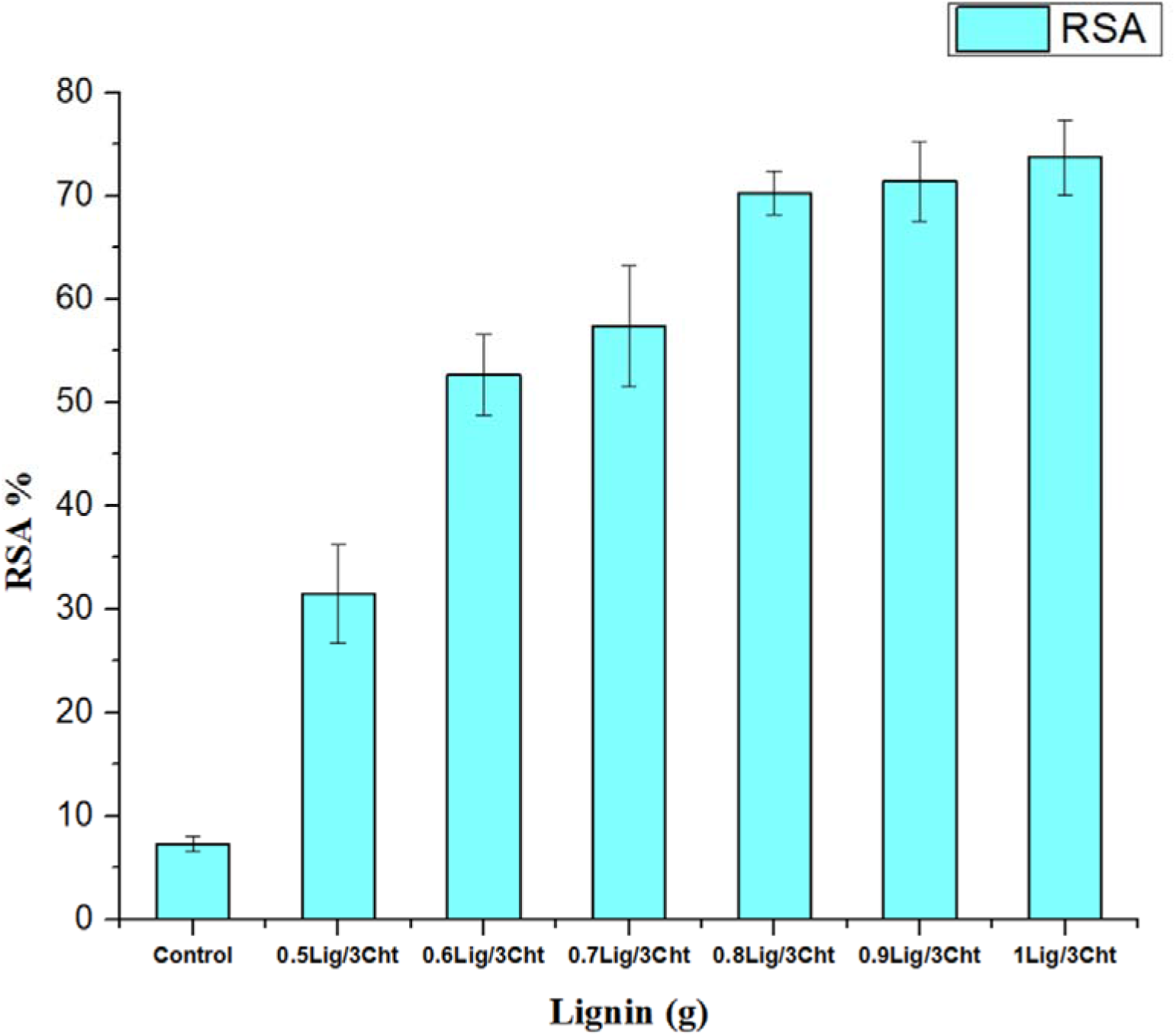
Radical scavenging activity of Lig/Cht films by DPPH assay.

#### 3.3.4 Water Vapor Transmissibility Rate (WVTR)

The water vapor transmissibility results reveal interesting trends amongst the chitosan hydrogels with varying lignin concentrations. Notably, as the lignin concentration increases from 0.6 to 0.8, there is a gradual decrease in water vapor transmission rates, with lowest rate observed for the 0.8Lig/3Cht hydrogel (66.22 ± 6.85 g/m²/h) as compared to 3Cht (control) (93.21 ± 0.22 g/m²/h). Reported studies on the inclusion of lignin in biopolymeric films resulted in a notable reduction in WVTR, decreasing to 50% of the control films [38]. However, further increment in lignin concentration to 0.9Lig/3Cht and 1Lig/3Cht did not follow this pattern (Fig. 6). This may be due to aggregation or other structural changes that could have compromised the mechanical integrity due to phase separation, causing the WVTR to increase [39]. This suggests that an optimal lignin concentration is essential to achieve the desired balance between water vapor permeability and mechanical strength of the Lig/Cht hydrogels which is in fact one of the reasons for incorporating lignin into the hydrogel as a reinforcing polymer material. Further investigation is warranted to understand the underlying mechanisms governing these observations and optimize the hydrogel formulations for specific applications. The current research can be useful in preserving foods that are prone to drying out or becoming stale such as baked goods such as bread, cakes, and pastries, as well as certain fruits and vegetables like berries, grapes, lettuce, and cucumbers. Additionally, with these hydrogels, other food materials such as meat, poultry, and seafood can also benefit by moisture retention and avoid drying out during storage or cooking. This packaging feature is particularly important in case of moisture containing food products as they profoundly degrade in terms of quality and texture on losing moisture. Therefore, implementing packaging solutions aimed at retaining moisture becomes vital to mitigate such deleterious effects and preserve the overall integrity of the food.

**Fig. 6.**
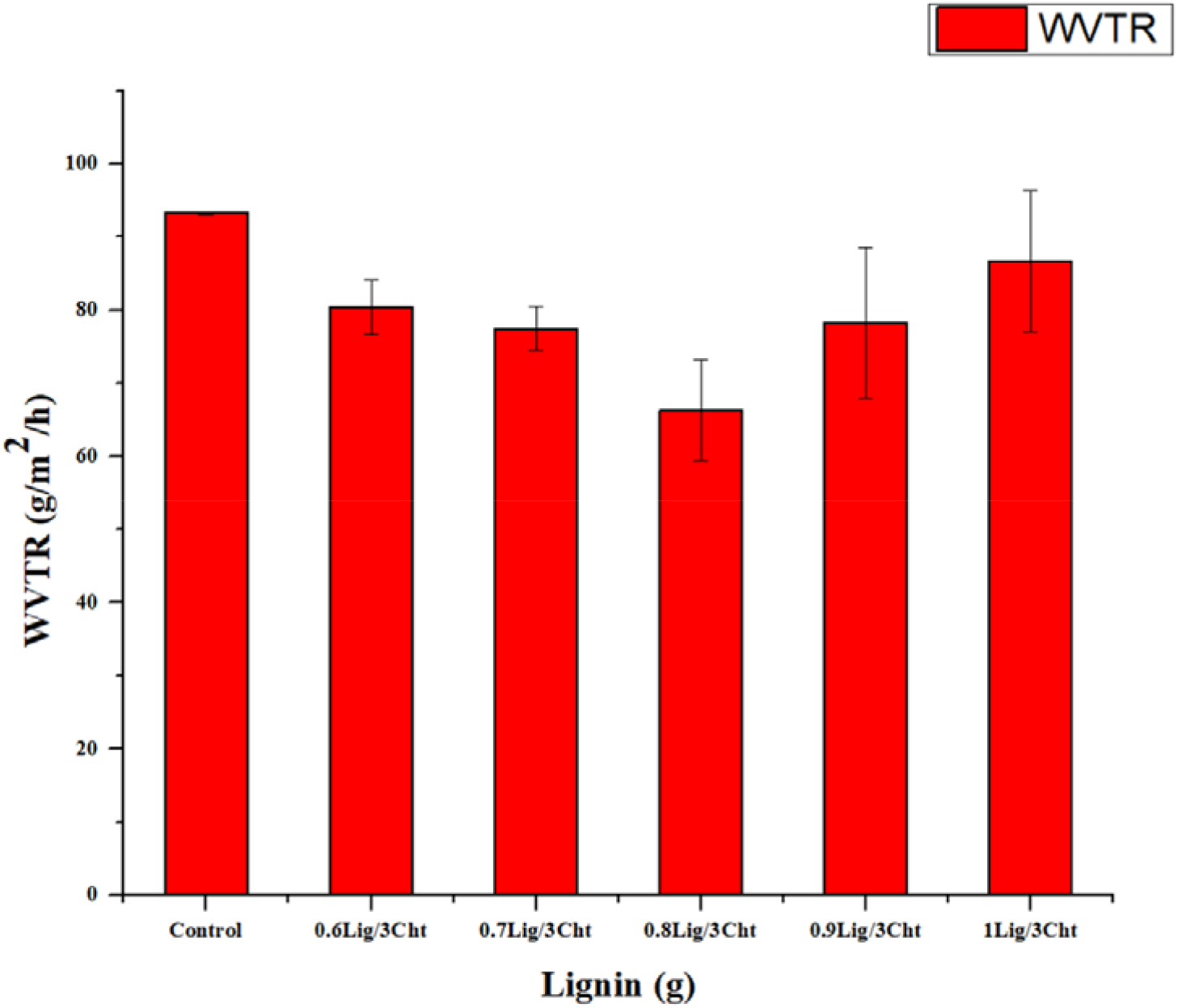
Profile of water vapour transmission rate (WVTR) by Lig/Cht and control (3Cht) films.

#### 3.3.5 Tensile strength

Tensile strength is determined by the cohesion between polymer chains, reflecting the material’s ability to withstand applied forces before rupture. On the other hand, elongation at the breaking point is indicative of the material’s flexibility and extensibility prior to failure [40]. The incorporation of lignin into hydrogels has been shown to enhance their mechanical properties, with improved stress transfer observed at the interface between the polymer matrix and lignin particles [41]. This phenomenon contributes to the reinforcement of the hydrogel structure, leading to enhanced tensile strength and Young’s modulus. The interaction between chitosan and lignin can be attributed to the formation of salt bridges between the protonated amino group of chitosan and the carboxylic group of lignin. These bonds facilitate the formation of a crosslinked framework within the hydrogel matrix, which bears the main mechanical load [42]. Hydrogels containing an optimal concentration of lignin fractions, namely, 0.8Lig/3Cht formulation, demonstrated improvement in both tensile strength and Young’s modulus. However, a visible decline in tensile strength is observed upon further addition of lignin beyond optimal value as shown in Fig. 7. This drop in mechanical properties may be attributed to possible irregularities in lignin dispersion within the hydrogel matrix, leading to weakened intermolecular interactions and decreased overall mechanical integrity [43]. Table 4 presents the various mechanical properties of chitosan hydrogels with different lignin concentrations.

**Fig. 7.**
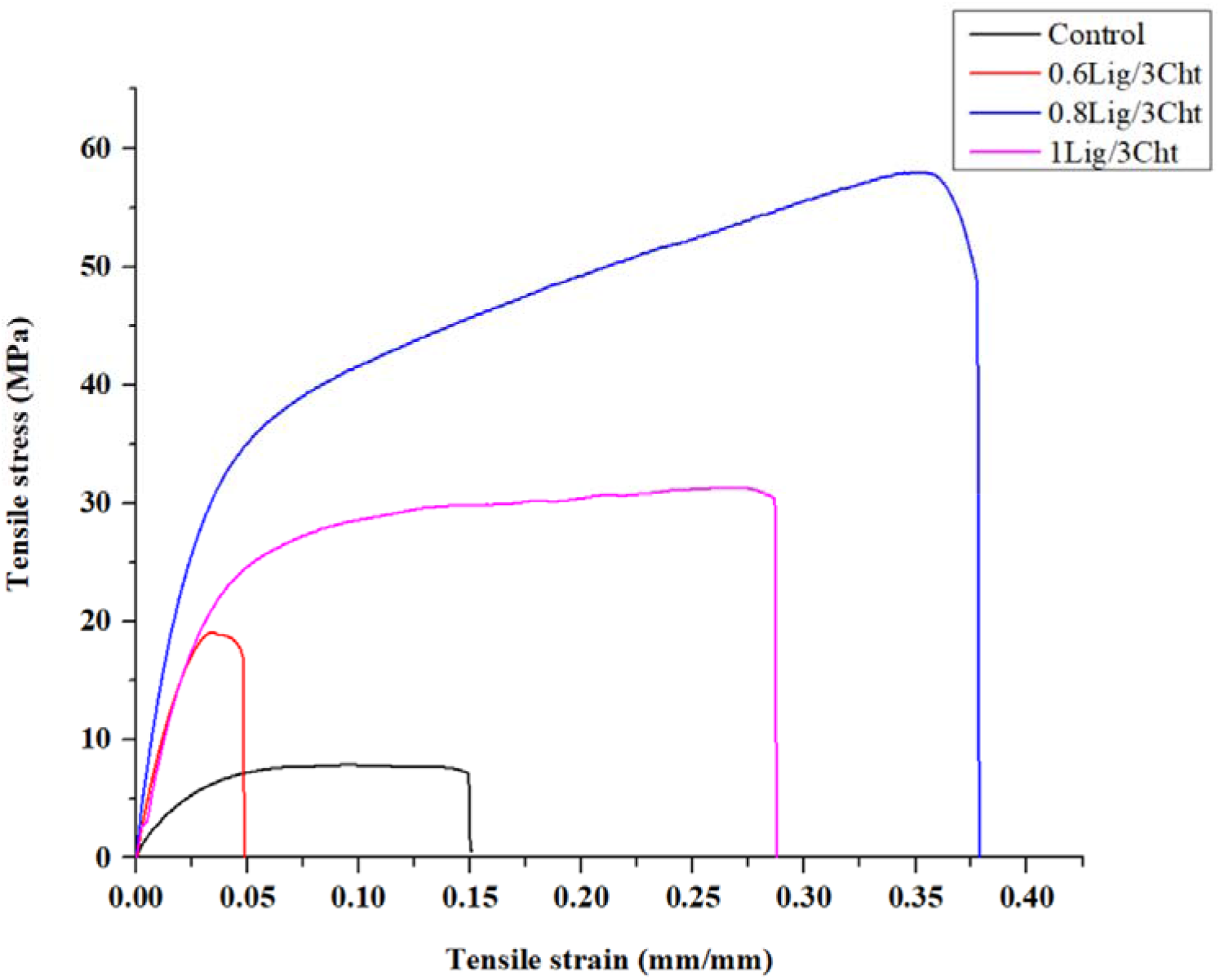
Stress-strain curve of Lig/Cht hydrogels and control (3Cht)

**Table 4.** Mechanical Properties of Chitosan Blend Hydrogels with Different Lignin Concentrations.

| Sample | Young's Modulus<br>[Mpa] | Load at<br>Break<br>(Standard)<br>[N] | Maximum<br>Load<br>[N] | Tensile<br>stress at<br>Break<br>(Standard)<br>[MPa] |
| --- | --- | --- | --- | --- |
| Control | 165.5 | 0.14 | 20.54 | 0.05 |
| 0.6Lig/3Cht | 627.22 | 35.43 | 38.19 | 17.71 |
| 0.8Lig/3Cht | 1037.42 | 49.29 | 57.95 | 49.29 |
| 1Lig/3Cht | 687.87 | 60.98 | 62.66 | 30.49 |

#### 3.3.6 Antimicrobial Property

The antimicrobial activities of control (3Cht) and Lig/Cht hydrogels were investigated against *Lactobacillus sakei* and *Bacillus subtilis* by a standard method [44]. Lactic acid bacteria can cause spoilage in meat and dairy products through the gradual accumulation of volatile organic acids [45]. On the other hand, *Bacillus subtilis*, a ubiquitous soil bacterium, is often associated with food spoilage of dairy products [46]. The control hydrogel displayed inhibition zone of 10 ± 0.5 mm in case of *L. sakei* as shown in Fig. 8. On the other hand, with the increase in lignin concentration the zone of *L. sakei* inhibition increased from 12 mm in 0.6Lig/3Cht to 14 mm in 0.8Lig/3Cht, respectively. However, with further increase in lignin concentration, the zone of inhibition did not increase. The control hydrogel (3Cht) also exhibited small inhibition zone measuring 10 ± 0.5 mm, indicating inherent, though not prominent, antimicrobial activity which can be attributed to the amino functional groups that electrostatically interact with microbial membrane [47]. As lignin concentrations were augmented within the hydrogel formulations, a trend of escalating inhibition zone was observed till an optimum lignin value. This phenomenon aligns with previous findings attributing antimicrobial activity to lignin, particularly its phenolic hydroxyl groups which are supposed to exert antimicrobial effects by disrupting bacterial cell membranes and inhibiting enzyme activities crucial for microbial survival [48]. However, the no further increase in inhibition zones with increase in lignin concentration suggests a saturation effect wherein the antimicrobial activity reaches a maximum threshold beyond which additional lignin incorporation does not confer any significant enhancement in antimicrobial efficacy. This saturation phenomenon could be attributed to the formation of intermolecular complexes of lignin. On the other hand, Lig/Cht hydrogels did not demonstrate any clear zones of inhibition towards *B. subtilis*. Nonetheless, it was observed that these hydrogels effectively impeded the growth of *B. subtilis* over the surface area of the hydrogel.

**Fig. 8.**
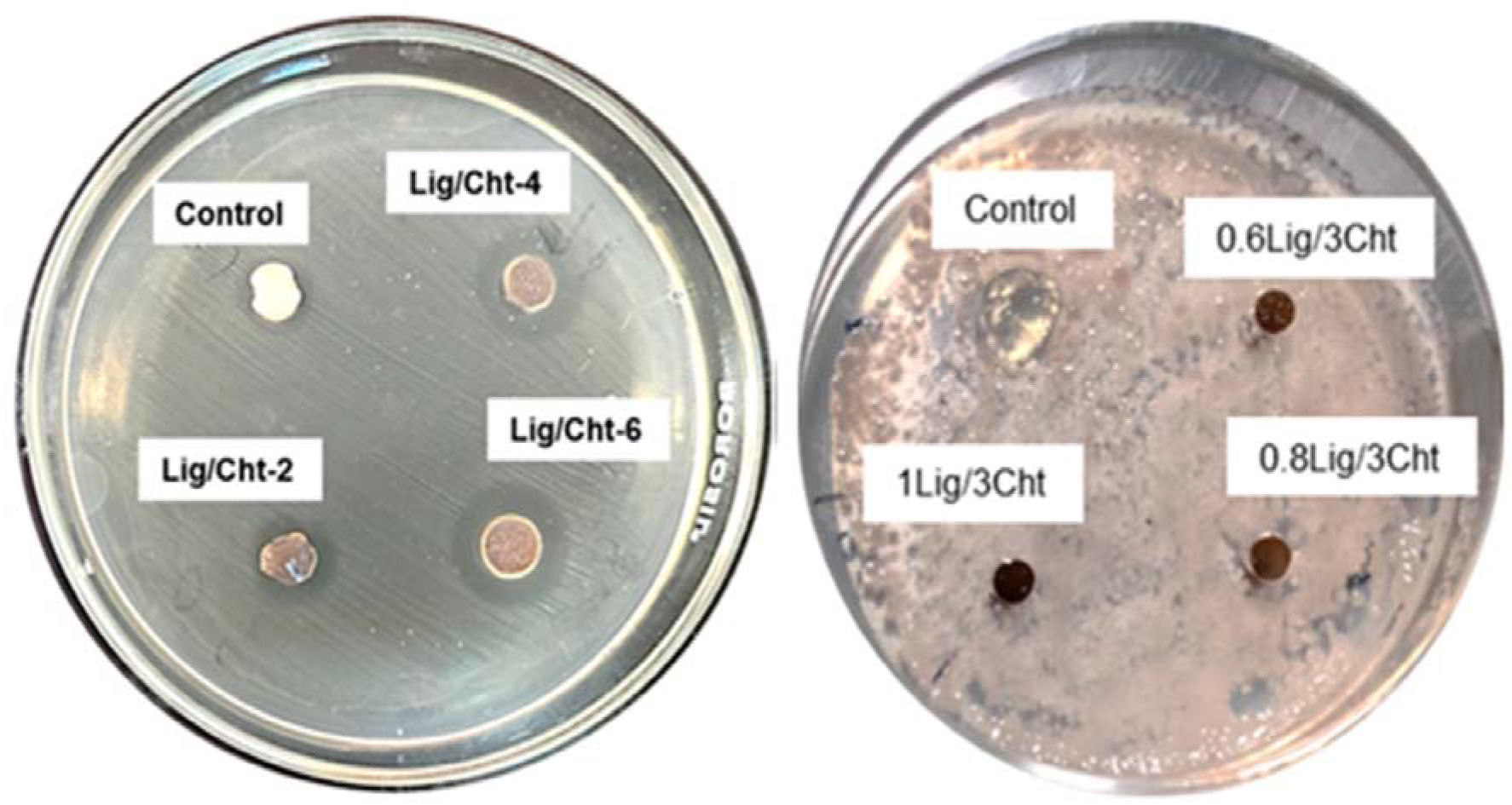
Antimicrobial activity of Lig/Cht hydrogels with **a.** *Lactobacillus sakei b. Bacillus subtilis*.

## Conclusions

The study presents a comprehensive investigation into the potential of lignin/chitosan hydrogels for food packaging applications. Comparison of FTIR characteristics between lignin extracted from sugarcane tops (SCTs) and commercial alkali lignin validates the successful lignin isolation process, paving the way for its utilization in hydrogel formulations. By varying lignin concentration, the impact on hydrogel properties was explored, including mechanical strength, UV barrier performance, antimicrobial, antioxidant, and water barrier properties. The findings underscore the importance of optimizing lignin content to achieve a balance between desired material properties, such as enhanced mechanical strength and UV blocking capabilities, while preserving antioxidant and antimicrobial efficacy. Further, the investigation into water vapour transmissibility rate (WVTR) revealed nuanced trends, emphasizing the need for careful optimization of lignin concentration to achieve its best performance in food packaging applications particularly for moisture rich products such as meat and dairy products. Hence, this research contributes to the growing body of knowledge on sustainable materials for packaging and underscores the potential of lignin/chitosan hydrogel as a versatile solution for improving food preservation through its reasonable bioactive properties. This study illustrates that utilizing lignin extracted from SCT in hydrogel formulations demonstrates a sustainable approach owing to the abundance of SCT as a renewable resource. Also, this study promotes exploitation of untapped resources and closely aligns with the principles of waste to wealth transformation, offering a promising avenue for sustainable green material development.

## Acknowledgements

Authors thank CSIR-ASPIRE (25WS[56216]/2023) for the research funding. SG acknowledges Council of Scientific and Industrial Research (CSIR), India for their financial assistance in the form of CSIR-NET Junior Research fellowship (09/1001(15595)/2022-EMR-1). Authors also thank the Director, Indian Institutes of Technology (IIT), Hyderabad for providing the seed grant (SG/IITH/F304/2022-23/SG-135) and other necessary facilities.

## Author Contributions

Ms. Sumona Garg: Writing – original draft, Data curation, methodology, Investigation.

Dr. Althuri Avanthi: Visualization, Supervision, Project administration; Resources, Writing - review & editing.

---

**Fig. S1.**
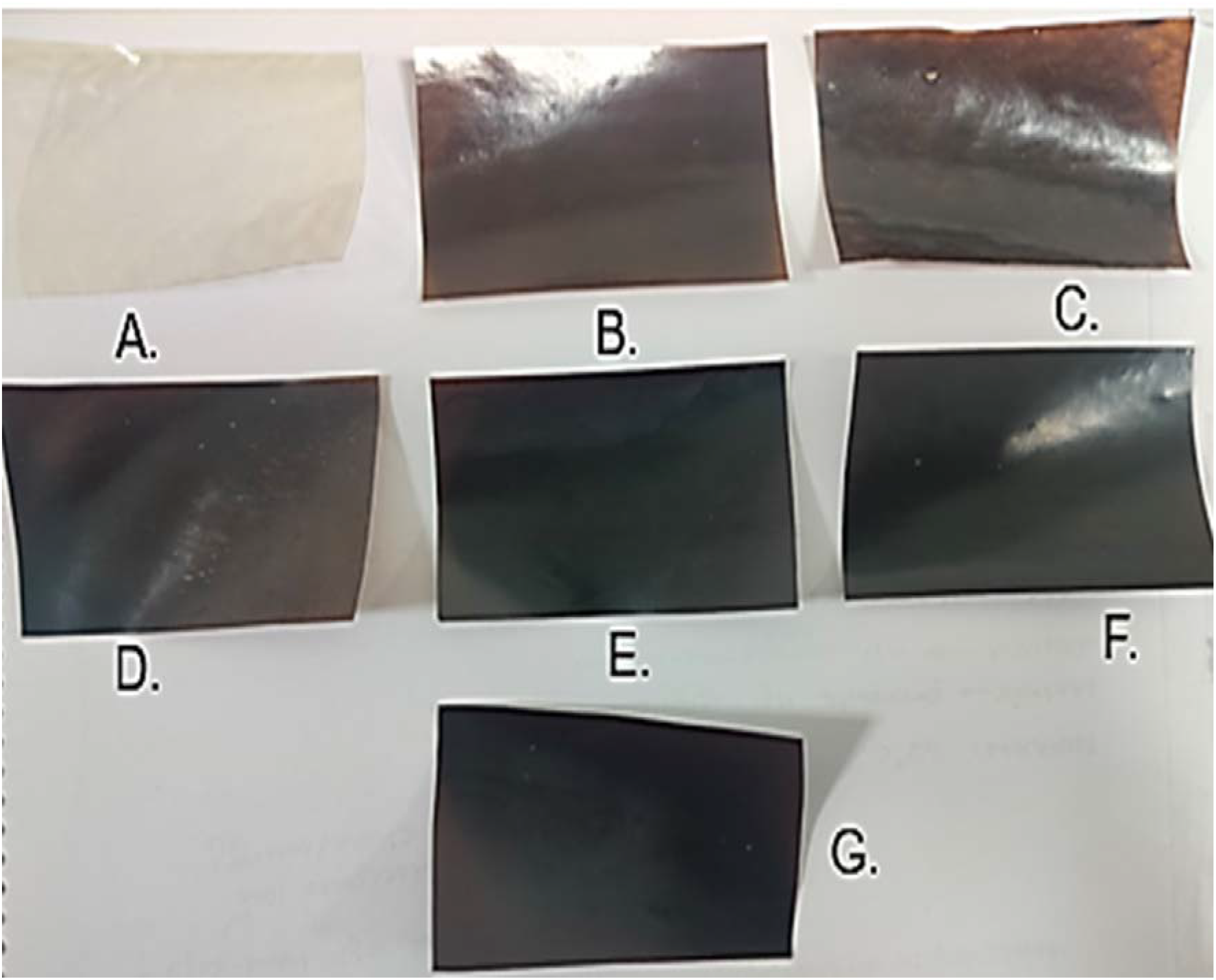
(A) Control (B) 0.5Lig/3Cht (C) 0.6Lig/3Cht (D) 0.7Lig/3Cht (E) 0.8Lig/3Cht (F) 0.9Lig/3Cht (G) 1Lig/3Cht.

